# Mistrafficking of KCC2 promotes hyperexcitability in hippocampal circuitry in a murine model of Christianson syndrome

**DOI:** 10.64898/2026.09.09.750201

**Authors:** Jamie Mustian, Haoyi Qiu, Andy YL Gao, Cong Loc Dang, Seyed Ehsan Vasegh, Bastien Castagner, John Orlowski, Reza Sharif-Naeini, R. Anne McKinney

## Abstract

How endosomal trafficking shapes inhibitory neurotransmission remains poorly understood, despite both processes being linked independently to epilepsy and neurodevelopmental disease. Christianson syndrome (CS), an X-linked neurodevelopmental disorder, represents a potential tractable condition to investigate such coupled processes. CS is caused by loss-of-function mutations in the *SLC9A6* gene which encodes the organellar (Na⁺, K^+^)/H⁺ exchanger NHE6 isoform whose loss overacidifies recycling endosomes and disrupts cargo delivery. Prior work has emphasized NHE6’s role in excitatory neurons and glia; whether endosomal dysfunction reshapes inhibitory circuits and thereby drives the early treatment-resistant epilepsy that defines CS has not been addressed. Using *Nhe6^-/Y^* mice, we identify a previously unrecognized endosomal-to-inhibitory axis. NHE6 is required for the surface delivery and stability of the neuron-specific potassium-chloride co-transporter KCC2 (encoded by *SLC12A5*) that sets the driving force for GABAergic inhibition. In hippocampal neurons, loss of NHE6 resulted in developmental downregulation and mistargeting of KCC2which impaired intracellular chloride homeostasis and enabled circuit-level hyperexcitability in response to a subthreshold convulsant challenge. These changes recast CS epilepsy not as a downstream consequence of excitatory dysfunction but as a primary failure of inhibitory development driven by endosomal mistrafficking of KCC2. More broadly, they establish endosomal pH regulation as a determinant of KCC2 biology, a node implicated across genetic and acquired epilepsies. Pharmacologically restoring KCC2 function therefore offers a mechanism-based therapeutic strategy for CS, and likely for the broader class of disorders in which endosomal trafficking and inhibitory imbalance converge.

**Statement of Significance:** Inhibitory neurotransmission depends on KCC2, a chloride extruder whose surface expression sets the polarity of GABA signaling. Yet how cells maintain KCC2 at the plasma membrane has remained unclear. We show that recycling endosomal pH, controlled in part by the (Na⁺, K^+^)/H⁺ exchanger NHE6, is a previously unrecognized determinant of KCC2 trafficking. This regulation fails in Christianson syndrome (CS), an X-linked disorder caused by loss-of-function mutations in NHE6. In CS, KCC2 is downregulated and mislocalized, chloride homeostasis collapses, and hippocampal circuits become hyperexcitable. The finding bridges two physiological processes, endosomal trafficking and inhibitory neuronal plasticity, and identifies a cellular checkpoint that could explain why endosomal disorders so often produce epilepsy. It also nominates KCC2-restoring therapeutics as a rational strategy for a currently untreatable disease.

## Introduction

Balanced excitatory and inhibitory (E/I) signaling is fundamental to brain function, and its disruption is a convergent feature of epileptic and many neurodevelopmental disorders[1]. While excitatory mechanisms have dominated investigations thus far, inhibitory failure is increasingly recognized as a primary driver of early-onset, pharmaco-resistant epilepsy[2, 3]. The strength and polarity of gamma-aminobutyric acid (GABA)-mediated (GABAergic) activation of its cognate receptors is set by the intracellular chloride (Cl^-^) gradient established by the opposing actions of two distinct alkali cation–Cl^-^ co-transporters; the Na^+^-K^+^-Cl^-^ cotransporter 1 (NKCC1) which imports Cl^-^ and dominates in immature neurons, and the K^+^-Cl^-^ cotransporter 2 (KCC2) which extrudes Cl^-^ and is progressively upregulated over the first postnatal weeks[4–6]. The developmental switch from NKCC1- to KCC2-dominant transport flips GABAₐ receptor activation from depolarizing (excitatory) to hyperpolarizing (inhibitory), a transition essential for the maturation of inhibitory circuits and for protecting the developing brain from hyperexcitability[7–9]. KCC2 surface expression is tightly governed by endocytic recycling, and even modest reductions in plasma membrane KCC2 collapse the Cl^-^ gradient and unmask network hyperexcitability[10, 11]. Yet the cellular machinery that controls KCC2 trafficking, particularly the endosomal sorting decisions that determine whether KCC2 is recycled to the surface or routed to lysosomes for degradation, remains incompletely defined.

Graded acidification of endocytic vesicles is a critical determinant of the processing and sorting of internalized cargo. The intraluminal pH of recycling endosomes is mildly acidic (typically ∼pH 6.5), driven by the vacuolar H^+^-translocating ATPase (V-ATPase) and maintained in part by the alkalinizing actions of the (Na⁺, K^+^)/H⁺ exchanger NHE6. Loss of NHE6 causes excessive acidification of recycling endosomes, accelerates cargo degradation, and disrupts surface delivery of multiple proteins essential for neuronal excitability and synaptic function[12–14]. NHE6 dysfunction has been studied largely in excitatory neurons and astrocytes, where it impairs brain-derived neurotrophic factor (BDNF)/tyrosine receptor kinase B (TrkB) signaling, dendritic arborization, and glutamatergic synapse maturation[12, 15, 16]. Whether dysregulation of recycling endosomal function also reshapes the inhibitory machinery, and KCC2 in particular, has not been addressed despite clinical observations that point strongly toward an inhibitory defect.

Such clinical indications come from Christianson syndrome (CS), an X-linked neurodevelopmental and neurodegenerative disorder caused by loss-of-function mutations in *SLC9A6*, the gene encoding NHE6[17, 18]. CS patients display multiple debilitating symptoms, including pronounced intellectual disability, microcephaly, ataxia, and autistic features with early-onset, drug-resistant epilepsy that typically presents within the first months of life[17–19]. Despite antiseizure medications, many affected boys die from sudden unexplained death in epilepsy (SUDEP) before age fifteen[19]. The disorder is among the more common forms of X-linked intellectual disability, with an estimated incidence of 1 in 16,000 to 1 in 100,000[19]. No disease-modifying therapy exists, in part because the mechanistic link between endosomal dysfunction and seizure generation has remained obscure. Notably, the early appearance of seizures, well before substantial neurodegeneration, and their onset within the developmental window of the NKCC1-to- KCC2 switch together suggest a primary failure of inhibitory maturation rather than progressive excitotoxic injury.

Here we examined whether NHE6 is required for the trafficking and function of KCC2, and whether its loss produces an inhibitory deficit sufficient to account for CS hyperexcitability. Using *Nhe6^-/Y^* mice[20], we show that hippocampal circuits are hyperresponsive to a subthreshold convulsant challenge, that KCC2 is developmentally downregulated and mislocalized in the absence of NHE6, and that Cl^-^ homeostasis is consequently impaired. These findings identify endosomal pH as an upstream regulator of inhibitory neurotransmission and reposition CS epilepsy as a primary consequence of KCC2 mistrafficking. This has direct therapeutic implications, given that KCC2-enhancing compounds are now in clinical development for other pathophysiologic conditions.

## Methods

### Transgenic animals

Adult wildtype C57BL/6 (Strain #:000664) and *Slc9a6/Nhe6* knockout (KO, *Nhe6^-/Y^*) (Strain #005843, name B6.129P2-*Slc9a6^tm1Dgen^*/J) mice were purchased from Jackson laboratories. The *Nhe6^-/Y^* mice were generated by targeted homologous recombination of a LacZ-Neo reporter cassette into an early coding region (exon six) of the *Slc9a6 gene* (disrupting normal transcription) in embryonic stem cells. The resultant mice were backcrossed a total of at least five generations. The B6.129P2-*Slc9a6^tm1Dgen^*/J mice were crossbred with our line of wild-type (WT) L15 mice expressing membrane-tethered GFP (mGFP) in a subset of CA1 hippocampal pyramidal neurons under transcriptional control of the Thy1 promoter[21]. For the parvalbumin (PV) interneuron electrophysiological recordings, a line of mice containing a floxed bright red fluorescent protein, tandem dimer Tomato (tdTom), was crossed with a genetically engineered mouse line that selectively expresses Cre recombinase in cells containing PV (PV-Cre)) to generate homozygous TdTom/PV-Cre mice. TdTom/PV-Cre were then crossed with *Nhe6* KO mice to generate TdTom/PV-Cre/N6. These mice provided endogenous labeling of PV interneurons. Only male mice were used in experiments due to the more pronounced phenotype reported in male patients with CS. All animal handling procedures were carried out according to the guidelines of the Canadian Council on Animal Care and the McGill University Comparative Medicine and Animal Resources.

### Immunoblotting

Mice were deeply anesthetized and decapitated before brain extractions. Hippocampi were sectioned and flash frozen in liquid nitrogen. Hippocampal tissue was homogenized with radioimmunoprecipitation assay (RIPA) buffer consisting of 1% Nonidet P-40, 0.5% sodium deoxycholate, 0.1% sodium dodecyl sulfate (SDS), 50 mM Tris-HCl (pH 8.0), 1.50 mM NaCl, 10 mM NaF, 1 mM sodium orthovanadate, 1 mM beta-glycerophosphate, and complete MINI protease inhibitors (Roche). Tissue lysates were sonicated, centrifuged at 13,000 revolutions per minute for 10 minutes at 4 °C, then the supernatant was removed. A bicinchoninic acid colorimetric assay (BCA) was performed on lysates to obtain baseline protein concentrations within each sample. For western blots, samples were prepared at a 1 µg/µl concentration and lysates were loaded in 8–9% SDS-polyacrylamide gel electrophoresis (SDS-PAGE), then transferred to methanol-activated polyvinylidene fluoride (PVDF) membranes (Millipore, Nepean, Ontario, Canada) overnight at 4 °C. Blocking of membranes was done using 5% non-fat skim milk or 5% bovine serum albumin/0.01% Tween-20 in phosphate-buffered saline (PBS-T) for 1 hour at room temperature, followed by incubation with primary antibodies diluted in blocking buffer. The following primary antibodies were used: NKCC1 (Abcam ab59791): 1:1,000; polyclonal anti-Rabbit KCC2 (Sigma 07-432): 1:1,000; monoclonal anti-mouse β-actin (Sigma A5441): 1:10,000; monoclonal anti-mouse β-III tubulin (Sigma T4026): 1:5,000. The following secondary antibodies were used: Peroxidase (HRP)- Affini Goat Anti-Rabbit (1:10000; Jackson 111-035-003) and Peroxidase (HRP)- Affini Goat Anti-Mouse (1:10000; Jacksin 115-036-062). The secondaries were incubated for 1 hour at room temperature followed by buffer washes. Then blots were imaged using the Amersham Imager600 and analyzed using ImageJ software.

### Immunofluorescence staining

Intracardiac perfusions and coronal sectioning were conducted in accordance with previously described methods. Adult male mice were deeply anesthetized then intracardially perfused with 0.1 M Phosphate-buffered saline (PBS) followed by 4% paraformaldehyde (PFA)/0.1 M PBS. Brains were post perfused 24-48 hours in PFA to ensure proper fixation, then submerged in 30% sucrose/0.1 M PBS until full saturation. Once ready, coronal sections of the brain were taken on 5100mz Vibrating Blade Tissue Slicer where each slice was 100 µm thick. Fixed slices were stored in 0.5% sodium azide/0.1 M PBS until subsequent immunolabeling. To begin immunolabeling, slice sections were permeabilized and incubated overnight at 4 °C in 0.4% Triton X- 100/1.5% HIHS/0.1 M PBS. After 24 hours, primary antibodies were diluted in permeabilizing buffer then slices were incubated on a shaker for 5 days in primary solutions at 4 °C. The following primary antibodies were used: polyclonal anti-rabbit NHE6 [13]: 1:500; monoclonal anti-mouse KCC2 (MA527510): 1:250; polyclonal anti-rabbit KCC2 (Sigma 07-432): 1:250; monoclonal anti-mouse EEA1 (Sigma E7659): 1:250; monoclonal anti-rat LAMP1 (DSHB 1D4B): 1:250; monoclonal anti-rat RAB11 (ab95375): 1:250; polyclonal anti-rabbit parvalbumin (Swant PV27): 1:500; and polyclonal anti-rabbit CAMK2α (ab131468): 1:250. After incubation and washing, secondary antibodies were diluted in 1.5% HIHS/0.1 M PBS and applied overnight at 4 °C. The following secondaries were used: Dylight 650 and Alexa Fluor 594. All slices were mounted onto SuperFrost (Menzel-Glaser) microscope slides using UltraMount fluorescence mounting medium (Dako) and 1.0 cover slips and left to dry overnight at room temperature in the dark then placed in 4°C cold room until ready to image.

### Antigen Retrieval Protocol

Slices stained with KCC2 (Sigma 07-432) underwent an antigen retrieval protocol before permeabilization to improve the penetration of the antibody. Coronal slices were submerged in a buffer containing 10 mM Tris/1 mM EDTA/0.05%Tween 20/ddH_2_O adjusted to pH 9.0 in a 24-well plate and placed in a bead bath at a temperature of 60 °C for 24 hours. The next day slices were allowed to cool to room temperature and washed with 0.1 M PBS. Then staining was proceeded as normal.

### Confocal Microscopy/Analysis

To capture high-resolution images of individual neurons, mounted slides were imaged using a Leica TCS SP8 confocal microscope with data acquired using a 63x (NA 1.4) HCXPL APO oil-immersion objective. mGFP and Alexa Fluor 488 were imaged using a 488 nm Ar laser line and Alexa Fluor 594 was imaged using a 543 nm He-Ne laser line. Channels were acquired separately to prevent spectral overlap of fluorophores. Optical sections of 300-500 nm were taken and line-averaged 2x at high resolution to improve the signal-to-noise ratio. To acquire whole brain sections, mounted slides were imaged on a Zeiss LSM800 Fully Automated Inverted Microscope using a 10X PLAN NEOFLUAR (NA = 0.30, Ph 1) objective. Tiling and z-stack were used to capture regions of interest. Scan times were determined automatically based on fluorescence intensity and minimized to prevent photobleaching. Tiles were then stitched together automatically into a composite image using ImarisStitcher software (Oxford Instruments). High-resolution confocal stacks were first deconvolved using Huygen Essentials software using a full maximum likelihood extrapolation algorithm (Scientific Volume Imaging, Hilversum, The Netherlands). 3D images were then compiled as maximum intensity projections using the Surpass function on Imaris software (Oxford Instruments). To quantify colocalization, the Surface function in IMARIS was used to create a mask of the tertiary dendrites using the mGFP channel. The channel containing signal for the protein of interest was masked based on the dendritic mGFP signal using the Imaris Surfaces function to ensure puncta were localized within neurons. For endosomal markers, the threshold for Surface creation was lowered such that the Surface was oversaturated relative to the mGFP signal in order capture puncta in direct apposition to the neuron of interest. The number and volume of puncta localized in the dendrite for each stack were then determined automatically using the Imaris Spots function. To quantify cell counts of CAMK2α+ neurons and PV+ interneurons, the Imaris Measurement Points function was utilized to measure the length (in μm) of the hippocampi regions. Manual counting of the neurons was done by three blinded experimenters. The number of neurons was normalized by dividing the length of the region by the total number of neurons to find the ratio of neurons per length (in μm).

### Synthesis of 6-Methoxy-*N*-ethyl-1,2-dihydroquinoline (DiH-MEQ)

#### SMILES: COC1=CC=C2C(C=CCN2CC)=C1

DiH-MEQ was prepared according to an established protocol[22]. Sodium borohydride (1.2 mg, 32 μmol) was added to a stirred solution of 6-methoxy-*N*-ethylquinolinium iodide (MEQ) (5 mg, 16 μmol) in distilled water (0.2 mL) under a nitrogen atmosphere. Upon addition, the stirred solution first turned red, then yellow. The solution was stirred at room temperature for 30 min. Thereafter, the solution was diluted with distilled water (3 mL). The aqueous layer was extracted with EtOAc (3 x 5 mL). Organic layers were combined, washed with brine (5 mL), dried with MgSO_4_, filtered and concentrated *in vacuo* to give DiH-MEQ as a red oil (3 mg, 16 μmol). DiH-MEQ was stored under a nitrogen atmosphere and used in the next step without further purification. DiH-MEQ was stable under a nitrogen atmosphere for at least two weeks at −80 °C and one week at −20 °C.

### Whole-cell patch recordings

Coronal brain slices were obtained from adult male mice (8 weeks old). Animals were deeply anesthetized through an intraperitoneal injection of 2,2,2-Tribromoethanol (Avertin, 250 mg/kg). Mice were then briefly perfused transcardially with an ice-cold oxygenated (95% O_2_, 5% CO_2_) N-Methyl-D-Glucamine based artificial cerebrospinal fluid (NMDG-ACSF) solution (bubbled with 95% O_2_ and 5% CO_2_) containing the following (in mM): 93 NMDG, 2.5 KCl, 1.25 NaH_2_PO_4_, 30 NaHCO_3_, 20 HEPES, 25 glucose, 2 thiourea, 5 Na-L-ascorbate, 3 Na-pyruvate, 12 N-acetyl-L-cysteine, 0.5 CaCl_2_/2 H_2_O, and 10 MgSO_4_/7 H_2_O (pH 7.3–7.4 adjusted with HCl 12 M). The brain was quickly excised and submerged in NMDG-ACSF. For brain slices, 400 µm transverse slices were cut with a vibratome (Leica VT1000S). Slices were transferred to a submerged chamber containing HEPES-based recovery ACSF for 10 minutes at 34 °C, equilibrated with 95% O_2_ and 5% CO_2_. Following the recovery incubation, slices were transferred to a recording chamber and continuously superfused with oxygenated ACSF (in mM): 119 NaCl, 24 NaHCO_3_, 2.5 KCl, 1.25 NaH_2_PO_4_, 2 CaCl_2_, 2 MgCl_2_, and 12.5 glucose (bubbled with 95% O_2_ and 5% CO_2_; pH 7.3; 300 ± 5 mOsm measured), where they were then maintained at room temperature prior to transfer to the recording chamber.

### Targeted whole-cell patch clamp

Slices were transferred to a recording chamber and continuously superfused with oxygenated ACSF (2 mL/min). Patch pipettes were pulled from borosilicate glass capillaries (Harvard Apparatus) with a P-97 puller (Sutter Instruments). They were filled with a solution containing (in mM) 135 K-Gluconate, 6 NaCl, 2 MgCl_2_, 10 HEPES, 0.1 EGTA, 2 MgATP, 0.8 NaGTP, 2 BAPTA (pH 7.3-7.4, adjusted with KOH; osmolarity, 300 mOsm, adjusted with sucrose) and had final tip resistances of 4–6 MΩ for whole-cell recording. Neurons in CA1 were viewed by an upright microscope (Olympus) with a 40X water-immersion objective, infrared differential interference contrast (IR-DIC) and fluorescence. Recordings were made in whole cell current clamp (holding potential at -70 mV) from identified PVNs expressing tdTom or pyramidal cell expressing mGFP. Data were acquired with pClamp 10.0 software (Molecular Devices) using MultiClamp 700B patchclamp amplifier and Digidata 1440A (Molecular Devices). Recordings were low pass filtered on-line at 2 kHz, digitized at 10 kHz and stored on a PC using pClamp software (Molecular Devices). After obtaining the whole-cell recording configuration, access resistance and membrane capacitance were calculated based on the response of a 10-mV hyperpolarizing voltage step from a holding potential of –70 mV. Recordings were corrected for liquid junction potential. Pipette offset was zeroed before and after recordings, and the cell was excluded if a drift of more than 5 mV was noted. Access resistance of more than 35 MΩ also excluded the cell from analyses. Three standardized current-clamp protocols were applied per cell recording. A gap-free 90 s-long recording with no injection current (I = 0) was used to measure the resting membrane potential (mV). A 1 s-long step current injection from –200 to +450 pA (50 pA step increments) was used to measure the spike count and spiking duration. A 1 s-long ramp current injection of varied slopes from 50 to 400 pA/s was used to measure the spike threshold. All action potential properties and intrinsic excitability analysis were performed on MATLAB software with the toolbox, ElecFeX [23]. For the convulsant experiment, pyramidal cells were left at resting membrane potential (I=0) and increasing concentrations of 4-AP (Sigma-Aldrich, A78403) were applied to the bath to induce spontaneous firing.

### Miniature EPSC/IPSC

Acute transverse hippocampal brain slices were prepared from male *Nhe6* KO mice (P30- P49), with age- and sex-matched WT as controls. After mice were deeply anaesthetized and decapitated, their brains were quickly removed and sectioned into 250 or 400 μm thick brain slices using a vibrating microtome (VT1000S, Leica), the chamber of which was filled with ice-cold sucrose-based artificial cerebrospinal fluid (ACSF). Slices were then allowed to recover in regular ACSF containing (in mM): 24 NaCl, 5 KCl, 1.25 NaH_2_PO_4_, 2 MgSO_4_, 26 NaHCO_3_, 2 CaCl_2_, and 10 glucose saturated with 95% O_2_/5% CO_2_ (pH 7.3, 300 mOsm) at 32 °C for 1 h before being maintained at room temperature prior to experimentation. Sections were bubbled constantly with 95% O_2_/5% CO_2_ in all the described preparation, recovery, and recording solutions.

To isolate AMPAR-mediated miniature excitatory postsynaptic currents (mEPSCs), 250 μm thick acute slices were placed into the recording chamber of an upright microscope (BX51WI, Olympus, XLUMPlanF1 20x 0.95 NA water immersion objective) and perfused continuously with ACSF (as described above) supplemented with (in μM): 1 TTX, 25 CPP, 50 picrotoxin, and 5 CGP 55845. Patch clamp recordings were then performed under whole-cell conditions from visually identified hippocampal CA1 pyramidal neurons held at -60 mV with an Axopatch 400 amplifier (Molecular Devices, Sunnyvale, CA, USA) at room temperature (23-25 °C) using borosilicate patch pipettes (4-8 MΩ) filled with (in mM): 120 K-gluconate, 1 EGTA, 10 HEPES, 5 MgATP, 0.5 Na_2_GTP, 5 NaCl, 5 KCl, and 10 phosphocreatine K2 (pH 7.2-7.3 with KOH and 285-295 mOsm). To monitor access resistance, transient test pulses were applied consistently every 2 min throughout the duration of the recording. Access resistance typically fell within the range of 7-10 GΩ, and data was discarded if the access resistance deviated > 20% during the recording. After holding current was stabilized, data was acquired at a sampling frequency of 20 kHz and filtered 123 at 2 kHz for 15 min. All AMPAR-mEPSCs were identified offline through use of Mini Analysis Software (Synaptosoft, Decature, GA). Thresholding for mEPSC amplitude detection was set at eight times the root-mean-square value of a visually-determined event-free recording span, and 450 events per cell were analyzed and utilized to determine mean values for each cell. A maximum of two cells was obtained from each slice before being discarded.

To isolate GABAAR-mediated mIPSCs, 1 μM TTX was also included in the external solution. Whole-patch-clamp recordings were performed on visually identified hippocampal CA1 pyramidal neurons or fluorescent transfected primary neurons held at -60 mV with an Axopatch 400 amplifier (Molecular Devices, Sunnyvale, CA, USA) at room temperature (23-25 °C) using borosilicate patch pipettes (4-8 MΩ) filled with (in mM): 120 K gluconate, 1 EGTA, 10 HEPES, 5 MgATP, 0.5 Na_2_GTP, 5 NaCl, 5 KCl, and 10 phosphocreatine K2 (pH 7.2-7.3 adjusted with KOH and 285-295 mOsm) (for intrinsic firing) or 140 CsCl, 4 NaCl, 174 0.5 CaCl_2_, 10 HEPES, 5 EGTA, 2 qx-314, 2 Mg-ATP, 0.5 Na-GTP (pH 7.36 adjusted with CsOH and 290 mOsm). The E/I balance was calculated as mean inter-event intervals of mEPSCs/mean of inter-event intervals of mIPSCs.

### Statistical Analysis

Statistical analyses were performed using GraphPad Prism. The data are represented as the mean ± the standard error of the mean (S.E.M.) from Student’s *t*-test (parametric data) or Mann-Whitney test (non-parametric data), as indicated in figure legends. The two-way ANOVA test was used when comparing the means of three of more independent groups, with post-hoc tests indicated in the figure legends. Statistical significance was defined as *p* < 0.05.

## Results

### Hippocampal circuits in *Nhe6^-/Y^* mice are hyperresponsive to subthreshold 4- aminopyridine challenge

Accurate information processing in the hippocampus depends on a tightly balanced ratio of excitatory to inhibitory synaptic drive, and even modest shifts in this balance can lower the threshold for epileptiform activity[24–26]. We therefore explored whether loss of NHE6 perturbs E/I balance in the hippocampus and, if so, whether it predisposes the circuit to hyperexcitability.

To this end, miniature excitatory and inhibitory postsynaptic currents (mEPSCs and mIPSCs, respectively) were recorded from CA1 pyramidal neurons in acute hippocampal slices prepared from brain tissue of P30-P49 WT and KO (*Nhe6^-/Y^*) mice. The E/I ratio— calculated from event frequencies—was significantly elevated in KO neurons, indicating a shift toward net excitation (**Fig. 1A**). Notably, KO mice do not develop spontaneous seizures despite this imbalance, suggesting that the underlying network may be sensitized rather than overtly hyperexcitable, primed to develop into epileptiform activity when pushed.

**Figure 1:**
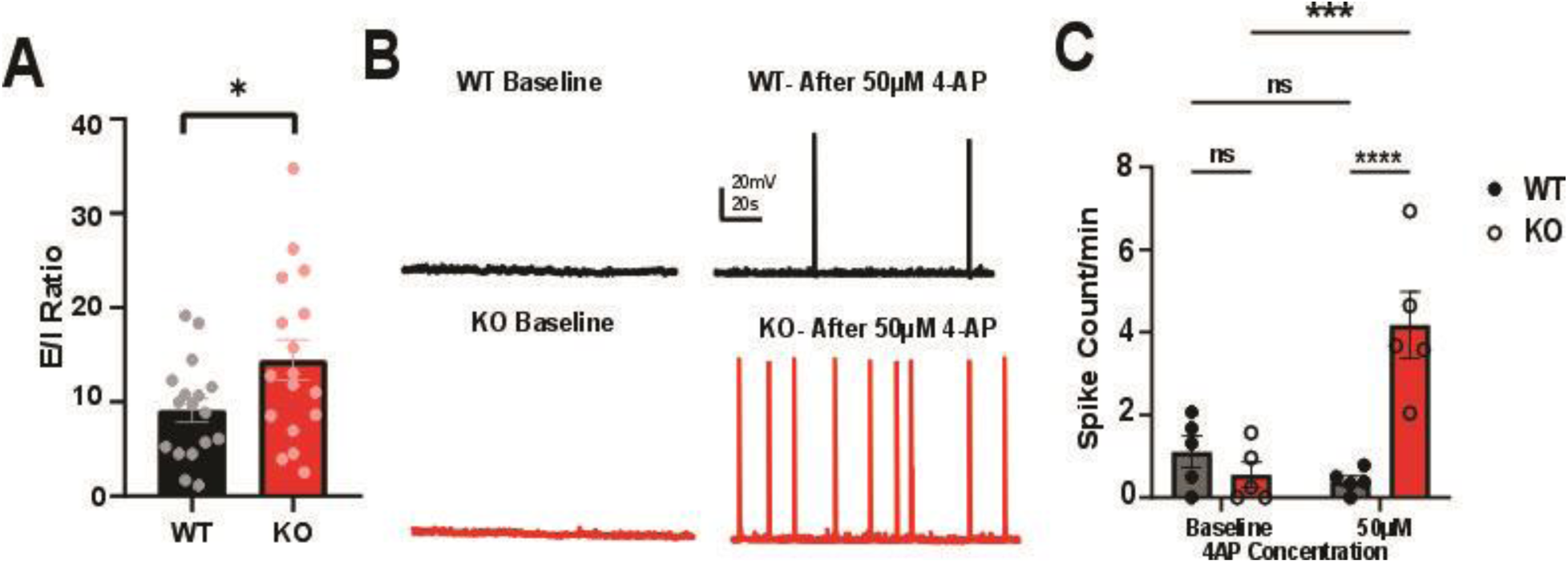
KO mice are more prone to hyperexcitability and demonstrate epileptiform activity compared to WT mice. A: Quantification of E/I ratio based on interevent intervals from mEPSC and mIPSC recordings in acute hippocampal slices. B: Example traces from whole-cell patch recordings of CA1 pyramidal neurons in gap-free of WT and KO acute hippocampi slices before and after 50 mM 4-AP bath application. C: Quantification of the spontaneous spikes per minute in B-C. WT: n = 5 cells from 3 mice, KO: n = 5 cells from 3 mice. *: p < 0.05; ***: p < 0.001, ****: p <0.0001, unpaired Student’s t-test, two-way ANOVA, Uncorrected Fisher’s LSD.

To test this directly, whole-cell current-clamp recordings were obtained from hippocampal CA1 pyramidal neurons in slices prepared from postnatal day 60 (P60) WT and KO mice, focusing on CA1 because of its well-documented vulnerability to seizure-induced injury[27]. Cells were held at their resting membrane potential while the K⁺- channel blocker 4-aminopyridine (4-AP) was bath-applied at sequentially increasing concentrations (20, 50, and 100M) to probe the network’s threshold for breakthrough firing. At baseline, spontaneous spike rates were low and comparable between genotypes (**Fig. 1B**, left; **Fig. 1C**; *supplemental Fig 2*). The KO CA1 pyramidal neurons showed a significant increase in spontaneous spike frequency at 50 and 100 μM 4-AP, whereas WT neurons remained unchanged across the full dose range (**Fig. 1B**, right; **Fig. 1C**). The representative traces capture the qualitative difference: WT cells remain quiescent under 4-AP, while KO cells generate repetitive suprathreshold events resembling early epileptiform discharges.

Together, these data show that CA1 circuits lacking NHE6 carry a constitutive E/I imbalance and, although silent at rest, break into epileptiform firing at 4-AP doses insufficient to perturb WT networks. This hyperexcitability mirrors the drug-resistant, threshold-lowered epilepsy that characterizes CS patients[28, 29] and prompted us to ask which synaptic component upstream of the imbalance is failing.

### Intrinsic firing properties of CA1 pyramidal neurons are altered in KO mice in the absence of changes in passive membrane properties

Having established that KO hippocampi are sensitized to convulsant challenge, we examined whether this network phenotype reflects a primary change in the excitatory cells themselves. NHE6 is expressed in the soma and dendrites of pyramidal neurons[13], raising the possibility that its loss alters either pyramidal cell number or their intrinsic excitability. Pyramidal cell density was quantified in adult WT and KO hippocampi using immunohistochemistry against the pyramidal marker CAMKIIα (**Fig. 2A**). CAMKIIα⁺ cell density was indistinguishable between genotypes (**Fig. 2B**), ruling out a gross loss of excitatory neurons as a contributor to the hyperexcitability described in **Fig. 1**.

**Figure 2:**
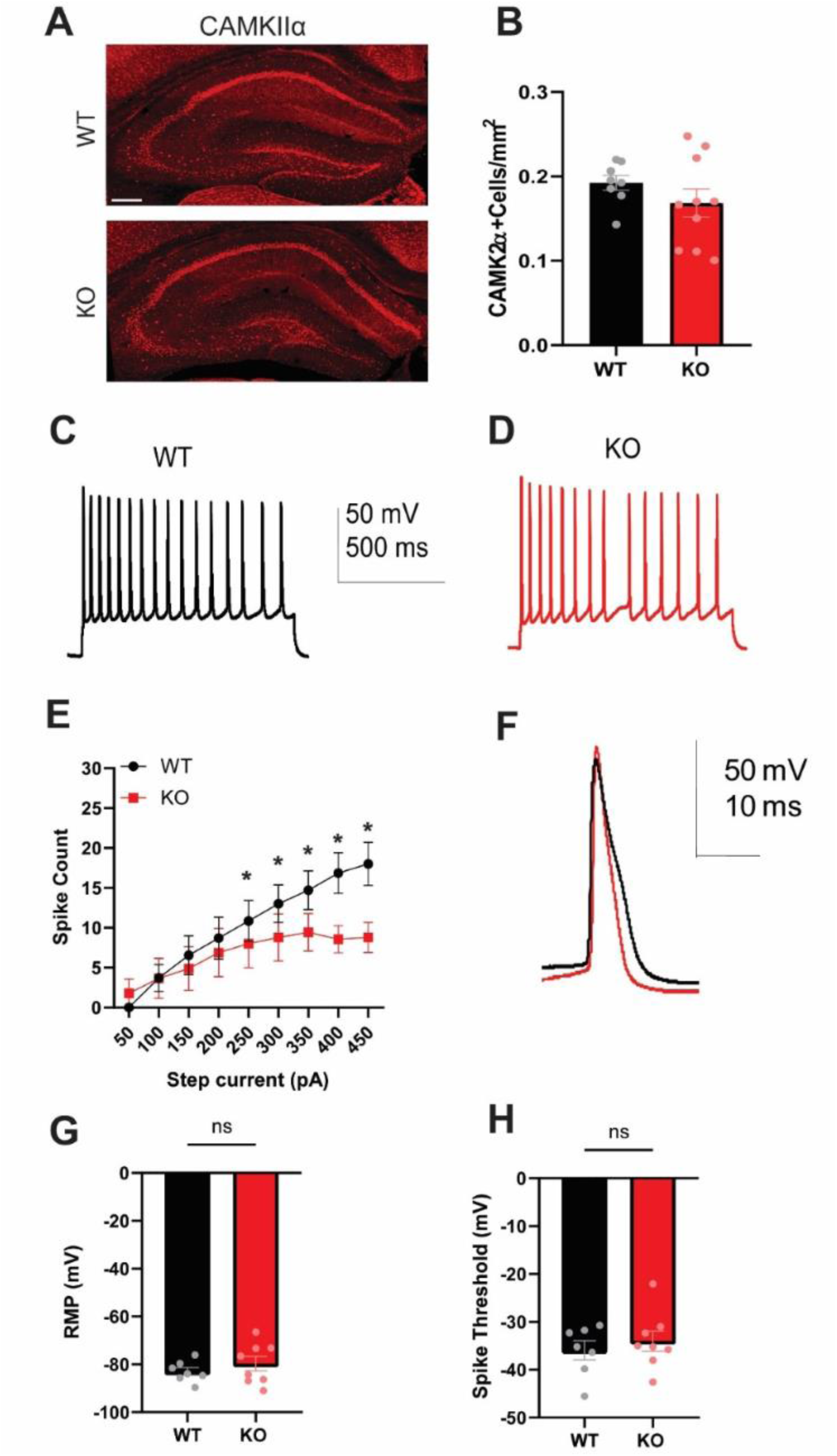
Intrinsic firing properties are altered in CA1 pyramidal neurons in KO mice in the absence of changes in passive membrane properties or cell density. A: Example immunofluorescent images of coronal brain sections taken from P60 CAMKIIα+ pyramidal neurons in WT and KO hippocampi (scale bar: 200 μm), B: Mean ± SEM quantification of CAMKIIα+ cell density. WT: n = 8 hippocampi in 4 sections from 3 animals; KO: n = 10 in 5 sections from 3 animals. C-D: Example traces from whole-cell patch recordings of CA1 pyramidal neurons following a current injection of +50pA, 1 sec in CA1 region of WT and KO acute hippocampi slice preparation. E: Spike Count. F: Inset of recordings for WT and KO. Mean ± SEM (F) resting membrane potential and (H) spike threshold. WT: n = 7 cells from 2 animals; KO: n = 8 cells from 4 animals. *: p < 0.05; **: p < 0.01, unpaired Student’s t-test and two-way ANOVA, Šídák’s multiple comparisons test.

Next, the intrinsic firing properties of CA1 pyramidal neurons were assessed by injecting a series of depolarizing current steps (50 to 450 pA) under whole-cell current clamp conditions. At low to moderate inputs (up to 200 pA), firing frequency was comparable between WT and KO neurons (**Fig. 2C, D, E**). At higher inputs (250 to 450 pA), however, KO neurons fired significantly fewer action potentials than WT, indicating a reduced firing capacity under a strong depolarizing current (**Fig. 2E**). Closer inspection of action potential waveforms during the 250-pA step revealed that individual spikes in KO neurons had a shorter half-width, a shorter fall time, and a faster fall rate compared with WT (**Fig. 2F**; *quantified in Supplementary Table 1*), pointing toward a subtle change in the repolarizing currents that shape the action potential. Passive membrane properties, including resting membrane potential (**Fig. 2G**) and spike threshold (**Fig. 2H**), were unchanged between genotypes.

Taken together, these data indicate that loss of NHE6 produces measurable changes in action potential waveform without rendering CA1 pyramidal neurons intrinsically more excitable; if anything, their firing output is reduced under strong depolarizing input. The hippocampal hyperexcitability documented in Fig. 1 therefore cannot be explained by a cell-autonomous increase in pyramidal neuron excitability and is more consistent with a deficit in the synaptic machinery that constrains these cells. These findings suggested that inhibitory interneurons might shape CA1 pyramidal output.

### Parvalbumin interneurons in KO mice show increased firing output without loss of cell density

Given that hyperexcitability of KO hippocampi did not arise from a cell-autonomous increase in pyramidal neuron firing, we next postulated that dysregulation of inhibitory parvalbumin-expressing (PV) interneurons might underlie this phenomenon because of their dominant role in pacing pyramidal output and the well-established link between PV dysfunction and epilepsy[30, 31]. This impetus was bolstered by analysis of RNAseq data in the publicly available DropViz adult mouse brain atlas (http://dropviz.org/) showing that cellular expression of *Nhe6* mRNA in the hippocampus is highest in GAD65 (glutamic acid decarboxylase 65)- and PV-positive interneurons[32]. Robust NHE6 protein expression was confirmed by confocal imaging of CA1 sections from mice in which PV+ interneurons were genetically labeled with tdTomato and co-immunostained for NHE6. The analysis revealed robust punctate signals for NHE6 within PV somata, consistent with its known endosomal distribution (**Fig. 3A**). The presence of NHE6 in PV+ interneurons raised two possibilities: that its loss compromises cell survival, or that it alters its function. The first was addressed by quantifying PV+ cell density in adult P60 hippocampi (**Fig. 3B**). PV+ interneuron density was unchanged between genotypes (**Fig. 3C**), arguing against PV cell loss as the basis for the network phenotype.

**Figure 3:**
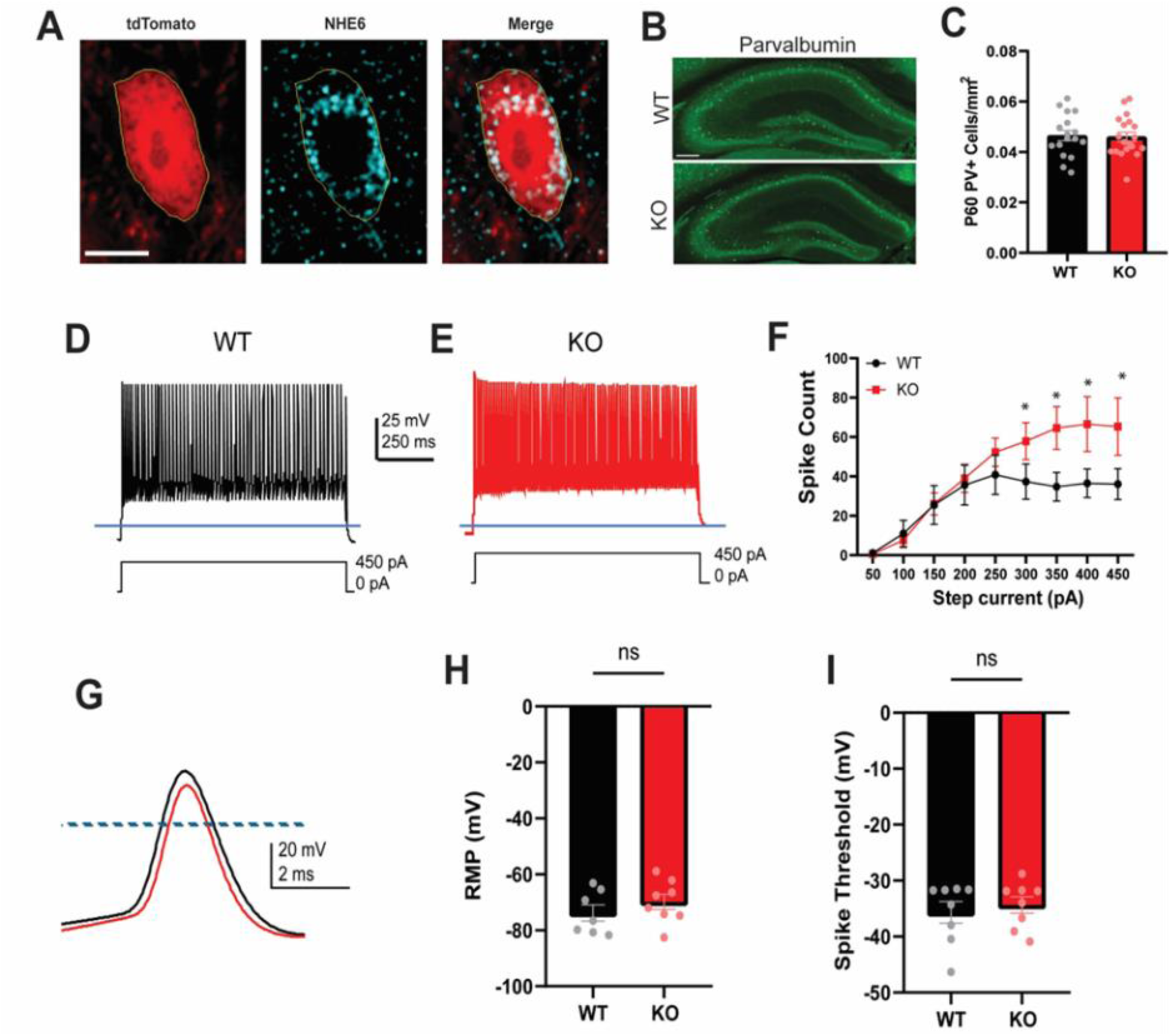
Spike count is elevated in KO PV interneurons despite no change in cell density and overall intrinsic firing properties. A: Example confocal images of tdTomato-labeled PV interneuron soma taken from a coronal CA1 hippocampal section from P60, immunolabelled for NHE6. Channels are shown separately and merged; the outline shows the soma. Scale bar = 20um. B: Example immunofluorescent images of coronal brain sections taken from P60 for parvalbumin in WT and KO hippocampi. (scale bar: 200 μm), C: Mean ± SEM quantification of PV+ interneuron counts. For PV, WT: n = 17 hippocampi in 10 sections from 5 animals; PV KO: n = 19 hippocampi in 10 sections from 6 animals. D-E: Example traces from whole-cell patch recordings of parvalbumin-expressing interneurons following a current injection of +50pA, 1 sec in CA1 region of WT and KO acute hippocampi slice preparation. F: Spike Count. G: Inset of recordings for WT and KO. Mean ± SEM resting (H) membrane potential and (I) spike threshold. WT: n = 7 cells from 2 animals; KO: n = 8 cells from 2 animals. *: p < 0.05; **: p < 0.01, unpaired Student’s t-test and 2-way ANOVA, Šídák’s multiple comparisons test.

The second possibility was examined by performing whole-cell current-clamp recordings of CA1 PV+ interneurons by injecting depolarizing current steps from 50 to 450 pA (**Fig. 3D, E**). In contrast to pyramidal neurons, KO PV+ interneurons fired significantly more action potentials than WT at high current injections (250 to 450 pA; **Fig. 3F**). Examination of the action potential waveform revealed a modest reduction in peak amplitude in KO PV+ cells (**Fig. 3G**; *quantified in Supplementary Table* 2), while resting membrane potential (**Fig. 3H**) and spike threshold (**Fig. 3I**) were indistinguishable between genotypes.

The directionality of this result is informative. Although GABAergic interneurons represent only a small fraction of hippocampal neurons, even subtle changes in their output reshape the timing and rhythmicity of pyramidal firing. Yet the increased spike output of KO PV+ interneurons cannot, on its own, account for the network hyperexcitability documented in Fig. 1. Instead, greater firing from PV+ interneurons would strengthen inhibitory drive, all else being equal. This pattern is more readily explained as a compensatory response, in which PV+ cells fire harder to offset a downstream failure of GABAergic transmission onto its pyramidal targets. We therefore investigated the postsynaptic determinants of inhibition; specifically, the developmental trajectory of the Cl^-^ homeostatic machinery that ultimately determines whether PV-driven GABA release is in fact inhibitory.

### Altered intracellular chloride homeostasis in KO hippocampi

The polarity of GABAergic signaling shifts during postnatal development from depolarizing to hyperpolarizing, driven by reciprocal regulation of two Cl^-^-coupled cotransporters: a developmental decrease in NKCC1, which loads Cl^-^ into the cell, and an increase in KCC2, which extrudes it[33]. Because NHE6 governs the endosomal trafficking and surface availability of multiple membrane proteins, its loss might perturb the developmental balance of NKCC1 and KCC2 in the hippocampus and thereby compromise mature inhibition. As a first step, expression of total NKCC1 and KCC2 protein was measured in whole hippocampal lysates from WT and KO mice at three postnatal time points (P21, P60, and P180) by western blot analyses. NKCC1 levels were comparable between genotypes at P21 but were significantly elevated in KO hippocampi at P60 and P180 (**Fig. 4A, C**). Conversely, KCC2 showed significantly reduced protein levels in KO hippocampi at P60 and P180 (**Fig. 4B, D**). The reciprocal changes produce a sustained imbalance in the Cl^-^ cotransporter ratio in adult KO hippocampus, one that intensifies with age and is consistent with a failure to fully execute the developmental NKCC1-to-KCC2 transition (*Supplemental Fig 3*). If this molecular signature has functional consequences, intracellular Cl^-^ should remain elevated in KO neurons.

**Figure 4:**
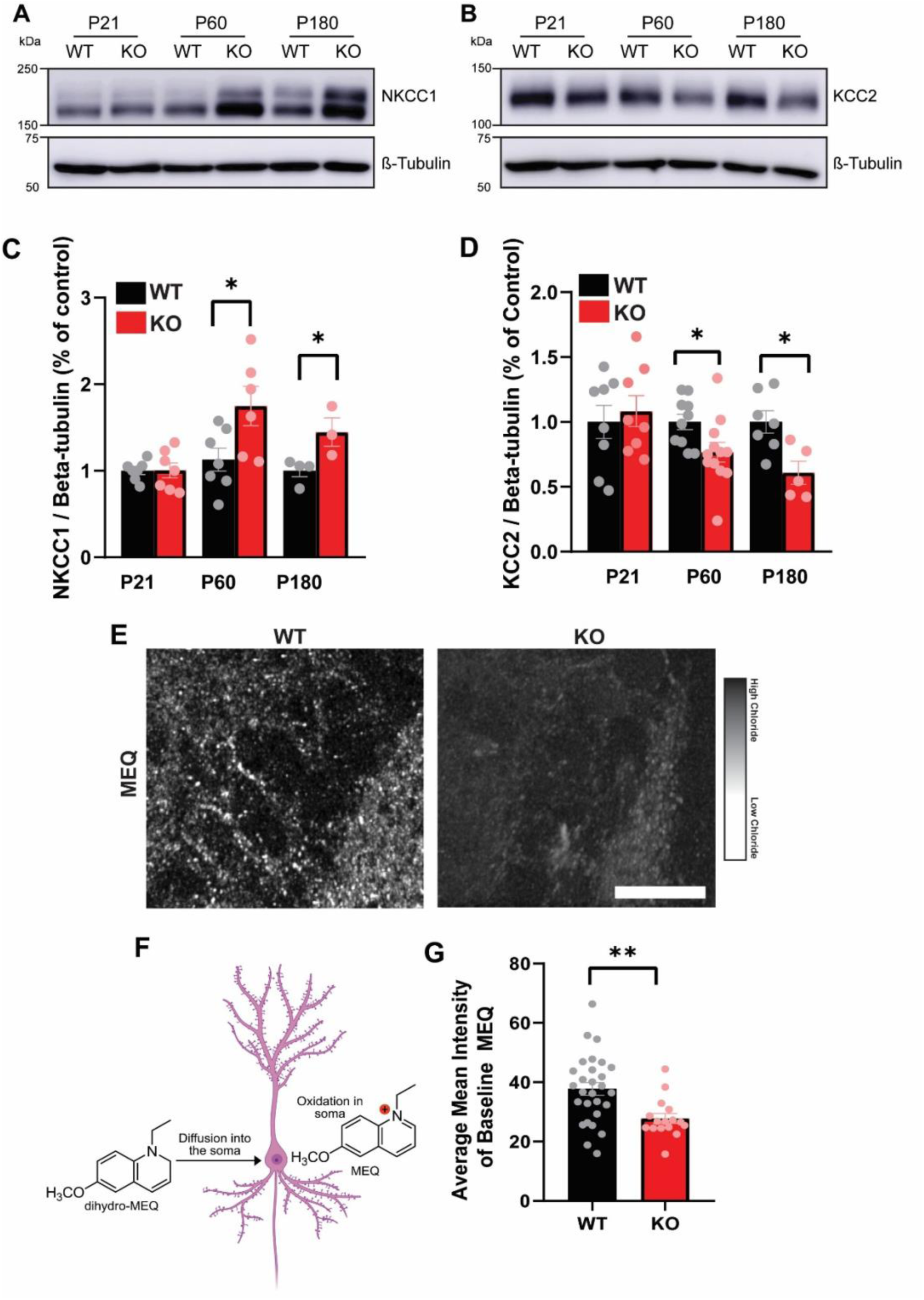
Upregulation of NKCC1 and downregulation of KCC2 in adult KO hippocampi and Living Imaging of Fluorescent chloride indicator (MEQ) in CA1 region. A-B: Example immunoblots of NKCC1 (A) and KCC2 (B) from P21, P60, and P180 WT and Nhe6 KO whole hippocampal lysates, with β-tubulin as loading control. B- C: Mean ± SEM quantification of NKCC1 (C) and KCC2 (D) relative to tubulin loading control and normalized to WT for each time point. E) Confocal images of adult hippocampal slices incubated in fluorescent dye (MEQ). F) Schematic of mechanism of entry and conversion of DiH-MEQ to MEQ in hippocampal neurons. G) Quantification of average MEQ fluorescence intensity. For NKCC1: P21, WT: n = 4, KO: n = 4; P60, WT: n = 7, KO: n = 8; P180: WT: n = 5, KO: n = 4. For KCC2: P21, WT: n = 6, KO: n = 6; P60, WT: n = 7, KO: n = 8; P180, WT: n = 5, KO: n = 4. *: p < 0.05, **: p < 0.01, unpaired Student’s t-test.

To test this prediction, intracellular Cl^-^ was imaged in acute hippocampal slices using MEQ, a fluorescent reporter whose emission is quenched in proportion to the intracellular Cl^-^ concentration; i.e., a brighter signal corresponds to lower Cl^-^[22]. Slices were loaded with the cell-permeant precursor dihydro-MEQ, which is oxidized within neurons to the membrane-impermeant, Cl^-^-sensitive MEQ and thereby retained intracellularly (**Fig. 4F**). The penetration of MEQ into slices was confirmed by incubating slices in 50 µM muscimol (a potent selective GABA_a_ receptor agonist) for WT or 10 µM bumetanide (an antagonist for NKCC1) for KO to visually reduce (increased Cl^-^) or elevate (reduced Cl^-^) fluorescence, respectively (*Supplemental Fig 4*). Live confocal imaging of the CA1 region in adult slices revealed significantly reduced MEQ fluorescence in KO compared with WT tissue (**Fig. 4E, G**), indicating elevated intracellular Cl^-^ levels consistent with the loss of KCC2 extrusion capacity. Collectively, these data show that loss of NHE6 disrupts the developmental regulation of Cl^-^ cotransporter expression in the hippocampus, producing a sustained Cl^-^ loading phenotype in adult CA1 neurons. A depolarizing shift in the GABA reversal potential is the expected consequence, offering a parsimonious explanation for the hippocampal network hypersensitivity. Specifically, GABA released by hyperactive KO PV+ interneurons would no longer exert its usual inhibitory effect on pyramidal targets, and the resulting collapse of network inhibition would render the circuit overreactive to convulsant challenge, as shown in Fig. 1.

### KCC2 is redirected to lysosomes in *Nhe6^⁻/Y^* hippocampi

The developmental failure of Cl^-^ homeostasis in *Nhe6^-/Y^* mice prompted us to ask how loss of NHE6 reduces KCC2 abundance. Previous studies have shown that KCC2 is internalized constitutively via clathrin-mediated endocytosis to the recycling endosomal compartment in heterologously expressing HEK293 cells[34], similar to NHE6[35]. This raises the possibility that NHE6 might colocalize with KCC2 and modulate its intracellular trafficking in hippocampal neurons. Consistent with this notion, confocal imaging of hippocampal dendrites immunolabeled for both KCC2 and NHE6 showed that a significant fraction of KCC2 overlapped with NHE6-positive endosomes (**Fig. 5A–C**), suggesting that NHE6 is positioned to potentially influence the membrane trafficking of KCC2.

**Figure 5.**
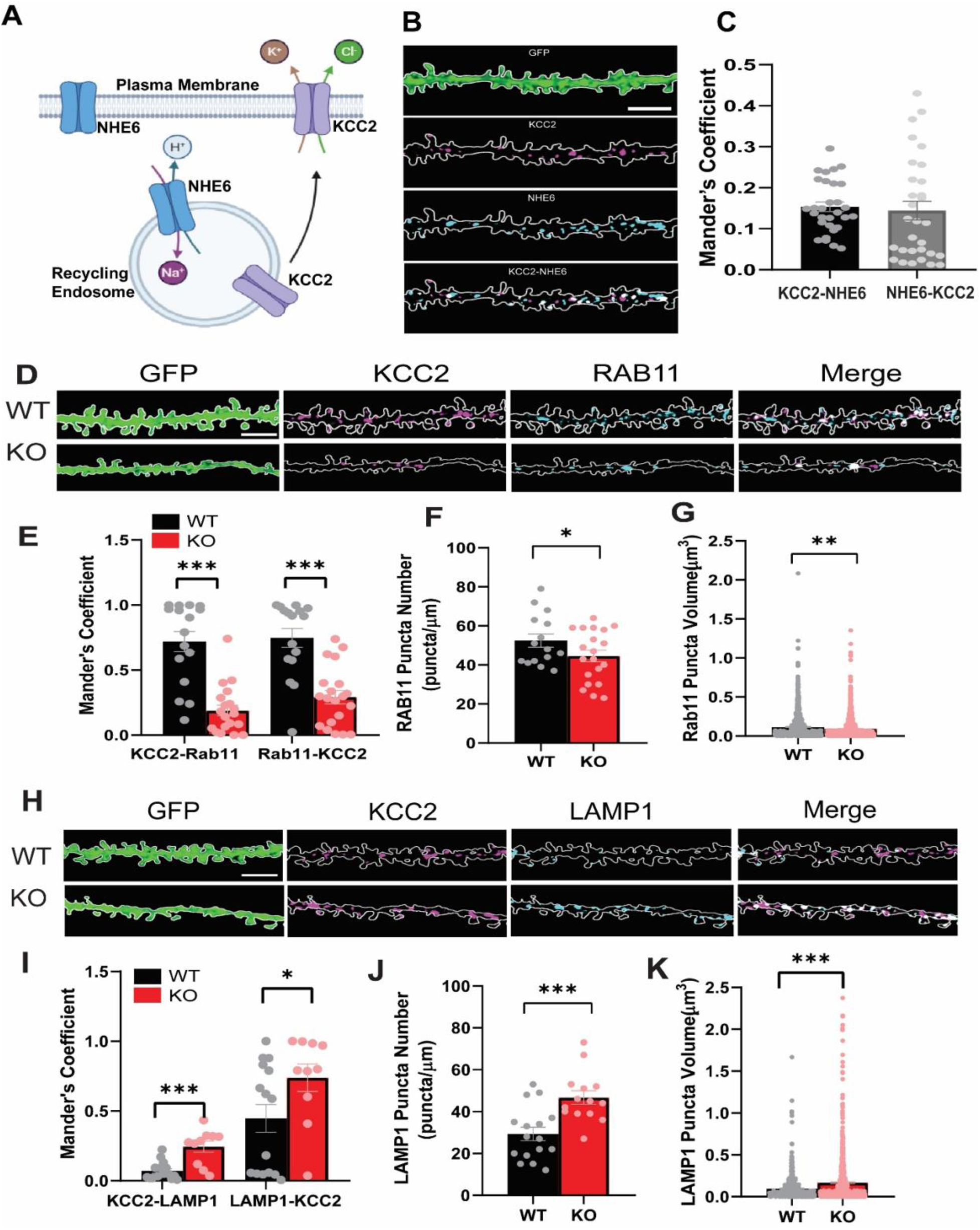
KCC2 is found in NHE6-containing endosomes and mistrafficked to lysosomes in KO. A: Schematic of KCC2 localization in mature WT neurons. B: Example confocal images of mGFP-labeled sections of tertiary dendrite taken from a coronal hippocampal section from P60 WT, immunolabelled with NHE6 and KCC2. Channels are shown separate and merged; dotted lines show outline of the dendrite denoted by the mGFP signal. C: Mean ± SEM quantification of Mander’s Coefficient for colocalization of KCC2 with NHE6. Scale bar: 4 μm. D: Example confocal images of mGFP-labeled sections of tertiary dendrite taken from a coronal hippocampal section from P60 WT and KO. Immunolabelled for KCC2 and RAB11. Channels are shown separate and merged; dotted lines show outline of the dendrite denoted by the mGFP signal. Mean ± SEM quantification of (E) Mander’s Coefficient for colocalization of KCC2 with endosomal markers, (F) Number of RAB11 puncta (G) Volume of RAB11 puncta. All quantifications represented as Mean ± SEM quantification and same information apply for H-K for LAMP1. For RAB11: WT: n = 16 dendrites in 8 hippocampi from 3 animals KO: n = 20 dendrites in 8 hippocampi from 3 animals. For LAMP1: WT n = 15 dendrites in 10 hippocampi from 3 animals KO = 13 dendrites in 10 hippocampi from 3 animals. Student’s t-test, *p<0.05, **p<0.01, ***p<0.001. Scale bar: 4 μm.

Because loss of NHE6 causes overacidification of recycling endosomes and promotes cargo mistargeting, we examined whether KCC2 trafficking is altered in the absence of NHE6. To this end, dual immunolabeling experiments were performed using markers of recycling endosomes (Rab11) and lysosomes (LAMP1) in hippocampal tissue from WT and KO mice. KCC2 colocalization with Rab11-positive recycling endosomes was significantly reduced in KO hippocampi relative to WT controls (**Fig. 5D–F**), indicating a loss of KCC2 association with the recycling endosomal pathway. In contrast, KCC2 colocalization with the lysosomal marker LAMP1 was significantly increased in KO hippocampi (**Fig. 5H–I**), suggesting that KCC2 is redirected away from recycling compartments to the degradative pathway.

If KCC2 is being rerouted from recycling endosomes to lysosomes, alterations in the morphology of these compartments would also be expected, as seen previously in hippocampal primary cultures of *Nhe6*-knockout neurons [12]. We therefore quantified both puncta number and compartment volume for Rab11-positive recycling endosomes and LAMP1-positive lysosomes. Consistent with impaired recycling, KO hippocampi exhibited a reduction in both Rab11 puncta number and recycling endosome volume compared with WT (**Fig. 5F–G**). Lysosomes showed the opposite pattern, with significant increases in both puncta number and volume in KO hippocampi (**Fig. 5J–K**).

Taken together, these findings identify KCC2 as cargo within NHE6-positive endosomal compartments and that loss of NHE6 shifts KCC2 trafficking away from the recycling pathway towards lysosomal compartments for degradation rather than for reuse at the plasma membrane.

## Discussion

Endosomal trafficking and inhibitory neurotransmission have largely been studied as distinct biological processes despite emerging evidence that they converge to shape neuronal circuit development and function. Here, we identify impaired NHE6-dependent trafficking of KCC2 as a mechanistic link between these pathways in CS. Although KCC2 dysfunction has been implicated in several epileptic disorders, it has generally thought to be a consequence of aberrant network activity. Although KCC2 dysfunction has been implicated in several epileptic disorders, genetic studies demonstrate that KCC2 dysfunction can also contribute directly to epilepsy. Rare variants of KCC2 have been identified in individuals with idiopathic generalized epilepsy and epilepsy of infancy with migrating focal seizures, with several variants producing impaired chloride extrusion, altered KCC2 surface expression or reduced transporter activity[36, 37]. Thus, KCC2 dysfunction can occur both upstream and downstream of epileptic activity. Our findings indicate that defective KCC2 trafficking, caused by loss of NHE6 function, contributes to altered hippocampal circuit excitability, and identify the recycling endosome as a compartment in which the fate of internalized KCC2 is determined. These results support KCC2 as a therapeutic target in CS and identify endosomal function as a determinant of inhibitory function.

Our results suggest that loss of NHE6 impairs recycling endosomal function and accelerates cargo degradation[12, 38]. We find that KCC2 resides within NHE6-positive compartments (**Fig. 5A–C**), placing it in a position to be affected by the loss of NHE6. In KO hippocampi, KCC2 is depleted from Rab11-positive recycling endosomes and enriched in LAMP1-positive lysosomes (**Fig. 5D–F, H–I**), and the compartments themselves shift in the same direction, with fewer and smaller recycling endosomes and more larger lysosomes (**Fig. 5F–G, J–K**). Consistent with a redistribution toward degradative compartments, total KCC2 protein is reduced, with a reciprocal elevation of NKCC1 (**Fig. 4A–D**), and intracellular Cl^-^ is correspondingly elevated (**Fig. 4E, G**). Importantly, this trafficking phenotype is present in animals that do not exhibit spontaneous seizures and is accompanied by reorganization of the recycling and lysosomal compartments themselves, indicating that the reduction in KCC2 reflects a primary sorting defect rather than a secondary consequence of seizure activity. GABA released onto CA1 pyramidal neurons would therefore be expected to exert a reduced hyperpolarizing effect. However KCC2 is expressed not only in pyramidal excitatory neurons but also in inhibitory neurons[39]. Therefore, the NHE6-depedent loss of KCC2 may affect chloride homeostasis across the hippocampal circuitry. The intrinsic properties we measured are consistent with a circuit responding to this loss. PV interneurons increase their spike output (**Fig. 3F**) while pyramidal neurons reduce theirs (**Fig. 2E**), two changes that in isolation would each oppose hyperexcitability. We interpret both as homeostatic responses to an upstream deficit arising from dysregulation of their Cl^-^ gradients. At rest these responses appear sufficient, and *Nhe6^-/Y^* mice do not develop spontaneous seizures; however, under subthreshold 4-AP challenge, KO circuits generate repetitive suprathreshold firing at concentrations that do not affect WT networks, revealing an underlying susceptibility to hyperexcitability in KO mice (**Fig. 1B, C**).

A key mechanistic question is how loss of NHE6 diverts KCC2 from recycling endosomes towards the degradation pathway. Two routes, one direct and the other indirect, are plausible. In the direct route, KCC2 is internalized from the plasma membrane by clathrin-mediated endocytosis and delivered to early/recycling endosomes[40, 41], where the recycling endosomal pool serves as a reservoir from which the surface complement is replenished³⁵. Our data on the colocalization place KCC2 in NHE6-containing endosomes (**Fig. 5A–C**). Upon loss of NHE6, there is a reduction of Rab11-labelled endosomes and a corresponding expansion of lysosomes (**Fig. 5F–G, J–K**), alterations consistent with a sorting defect rather than reduced synthesis of KCC2. In the indirect route, KCC2 is affected through other NHE6-dependent cargoes. BDNF/TrkB signaling stabilizes KCC2 at the plasma membrane in early development[42], and loss of NHE6 reduces TrkB availability[12]. The reduced TrkB signaling could therefore lower KCC2 stability. KCC2 surface stability is also regulated by phosphorylation of the c-terminus, specifically at the serine 940 residue. Phosphorylation of serin 940 by Protein kinase C reduces transport internalization and accumulation at the plasma membrane of KCC2, thus influencing chloride extrusion[41]. Conversely, dephosphorylation of serine 940 promotes KCC2 internalization and loss of KCC2 function has been associated with manifestation of seizure activity[10, 41]. Consistent with a contribution from impaired posttranslational regulation, we observe reduced phosphorylation at serine 940 in the CS model (*Supplementary Fig. 1*), as reported in other epilepsies[11, 43, 44].Our data favor the first route without excluding the second, and may involve both processes.

KCC2 membrane stability is additionally regulated by its mobility within the neuronal plasma membrane[45]. Rather than remaining stationary after insertion, KCC2 undergoes lateral diffusion between extrasynaptic regions of the plasma membrane and synaptic/perisynaptic microdomains at excitatory and inhibitory synapses, and increased mobility facilitates its removal from membrane clusters and subsequent endocytosis[46]. GABA_A_ receptor-mediated inhibition provides an activity-dependent mechanism for regulating this process: enhancing GABA_A_ receptor transmission restricts KCC2 lateral diffusion, increases its confinement at the plasma membrane and limits endocytosis, whereas antagonizing GABA_A_ receptor activity increases KCC2 membrane mobility and internalization effect[45, 47]. Thus, impaired inhibitory transmission in the CS hippocampus could further destabilize the remaining surface pool of KCC2, creating a feed-forward mechanism in which reduced KCC2 weakens GABAergic inhibition which, in turn, promotes further loss of surface KCC2. These routes are experimentally separable and distinguishing them by asking whether KCC2 mistrafficking persists when TrkB signaling is restored in *Nhe6^-/Y^* neurons is an important next step.

The developmental timing of the Cl^-^ transport phenotype differs from that predicted by a simple failure-of-maturation model. During normal postnatal development, the transition from NKCC1- to KCC2-dominant Cl^-^ transport converts GABA-_A_ receptor- mediated responses from depolarizing to hyperpolarizing, a process essential for the maturation of inhibitory neurotransmission[4, 48]. Impaired maturation of this developmental switch has been reported in several neurological disorders, such as chronic pain and epilepsy[49–52]. In contrast, KCC2 and NKCC1 expression did not differ between genotypes at P21, when the developmental Cl^-^ shift is largely complete in the mouse hippocampus, and diverged at P60 and P180 (**Fig. 4A–D**). These findings indicate that loss of NHE6 does not primarily impair establishment of the mature Cl^-^ transport profile but instead compromises its long-term maintenance, an interpretation consistent with the age-dependent worsening of the molecular phenotype. Defining the onset of this divergence at earlier postnatal ages will further resolve how endosomal trafficking contributes to the acquisition, as opposed to the maintenance, of mature Cl^-^ homeostasis.

Another observation of interest is the cell-type specificity of the intrinsic firing changes. Increased PV interneuron output alongside reduced pyramidal firing is consistent with compensation for GABAergic transmission that has become less effective at pyramidal targets. Alternative explanations include a cell-autonomous effect of NHE6 loss in PV interneurons, which express NHE6 (**Fig. 3A**), or reduced inhibitory drive onto PV cells. The accompanying change in pyramidal action potential waveform, a shortened half-width with faster fall time (**Fig. 2F**), indicates altered repolarizing conductance, consistent with NHE6 acting on additional membrane cargoes. KCC2 also has a structural role independent of Cl^-^ transport, contributing to dendritic spine maturation and excitatory synapse stability[53]. Notably, reduced dendritic branching and spine density have also been reported in NHE6-deficient neurons[12, 16]. Mistrafficking of KCC2 may therefore contribute to altered excitatory synaptic architecture alongside inhibitory dysfunction. The inhibitory network of CA1 itself is heterogeneous, and extending this analysis to somatostatin-expressing interneurons, as well as to the astrocytic compartment in which NHE6 is also expressed, will further define how endosomal trafficking shapes circuit excitability.

These findings have specific therapeutic implications. Strategies for enhancing KCC2 include small molecule compounds like CLP257 and its carbamate prodrug CLP290, which have been shown to restore Cl^-^ extrusion in neuropathic pain [54] and indirect approaches that increase KCC2 expression or stability through WNK–SPAK/OSR1, GSK3β or TrkB signaling[47, 55]. Because the KCC2 defect in CS is one of trafficking and stability rather than of intrinsic transport function, agents that protect KCC2 from lysosomal degradation, or that correct endosomal pH, may be suitable therapeutic strategies. Testing whether pharmacological enhancement of KCC2 normalizes network responses in *Nhe6^-/Y^*hippocampus is a direct extension of the present work. The reciprocal elevation of NKCC1 (**Fig. 4A, C**) also raises the possibility of inhibiting NKCC1, although the clinical record for bumetanide, an NKCC1 antagonist, has been mixed and its central nervous system penetration is limited[56, 57]. Our developmental data further suggest that because the deficit reflects impaired maintenance rather than impaired establishment of KCC2, the therapeutic window may extend beyond early development, a consideration of practical importance in a disorder typically diagnosed after seizure onset.

In summary, we identify NHE6 as a determinant of KCC2 stability and, through it, of inhibitory function in the hippocampus. We propose that the epilepsy of Christianson syndrome arises primarily from inhibitory rather than from excitatory dysfunction. More broadly, these results establish recycling endosomal function as a regulator of the Cl^-^ gradient on which inhibition depends and predict that other disorders of endosomal sorting should converge on impaired inhibition and increased seizure susceptibility. NHE6 function has been reported to be altered in Alzheimer’s and Parkinson’s disease[58, 59], and whether a comparable mechanism operates in those settings remains to be determined.

