## Supplementary Figures for "Mistrafficking of KCC2 promotes hyperexcitability in hippocampal circuitry in a murine model of Christianson syndrome"

**Supplemental Figures**

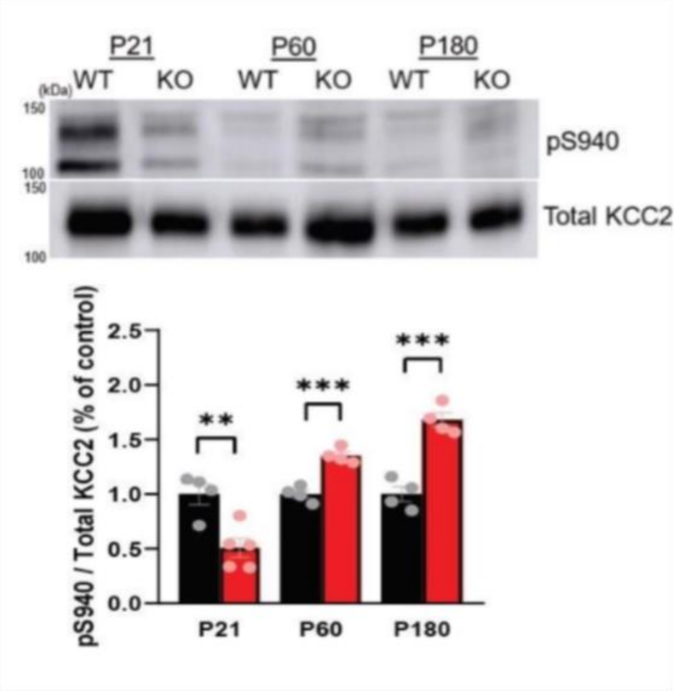


**Supplemental Figure 1:** ***Reduced activation of KCC2 in NHE6 KO***. *Example immunoblots of phosphorylated serin 940 (pS940) and total KCC2 from P21, P60, and P180 WT and KO whole hippocampal lysates, KCC2 as a control. Mean ± SEM quantification of pS940 relative to total KCC2 and normalized to WT for each time point. WT: n = 4, KO: n = 4-5. Student's unpaired t-test. *=p<0.05, **=p<0.01, ***=p<0.001.*

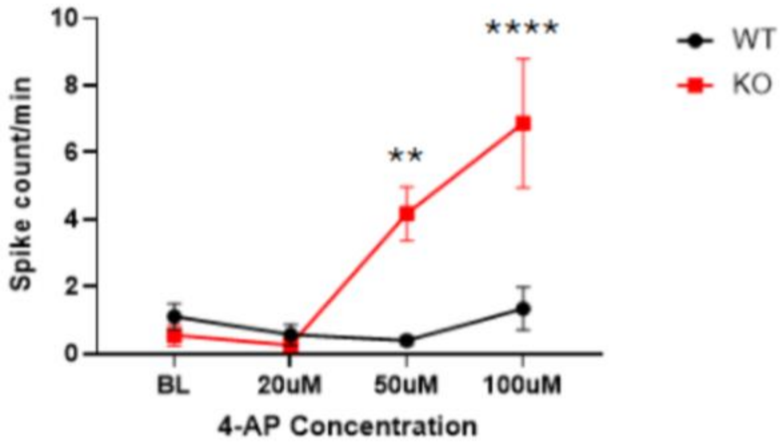


***Supplemental Figure 2: Quantification of spontaneous spikes per minute to increasing concentrations of 4-AP****. WT: n=5 cells from 3 mice, KO: n=5 cells from 3 mice. **: p < 0.01, ****: p <0.0001, unpaired Student's t-test, 2-way ANOVA, Tukey's multiple comparisons test.*

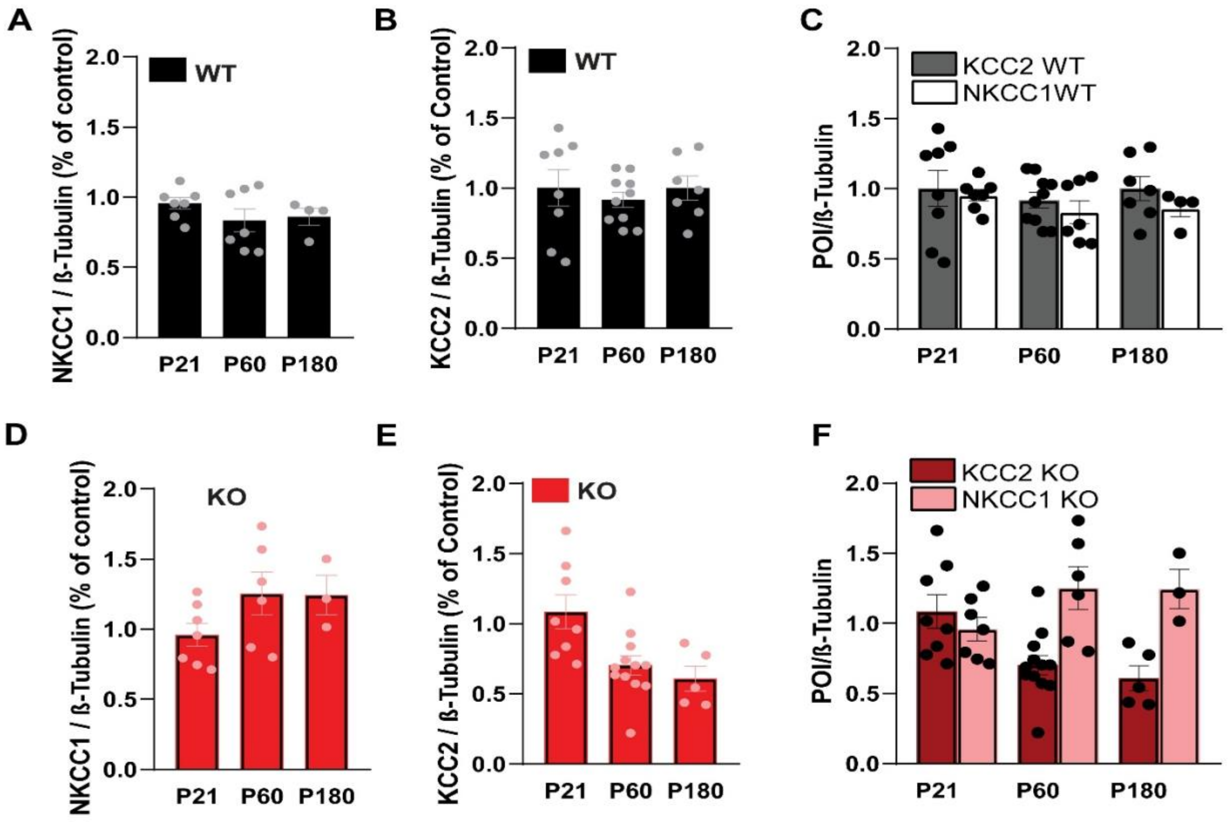


**Supplemental Figure 3: Time course of levels of KCC2 and NKCC1 in Adult Hippocampi**. *Mean ± SEM quantification of WT at P21, P60, and P180 for NKCC1 (A) and KCC2 (B) and combine (C) relative to tubulin loading control. Mean ± SEM quantification of KO at P21, P60, and P180 for NKCC1 (D) and KCC2 (E) and combine (F) relative to tubulin loading control. Data is not normalized to demonstrate developmental trends in NKCC1 and KCC2 in WT hippocampi. For NKCC1: P21, WT: n = 4, KO: n = 4; P60, WT: n = 7, KO: n = 8; P180: WT: n = 5, KO: n = 4. For KCC2: P21, WT: n = 6, KO: n = 6; P60, WT: n = 7, KO: n = 8; P180, WT: n = 5, KO: n = 4. unpaired Student's t-test.*

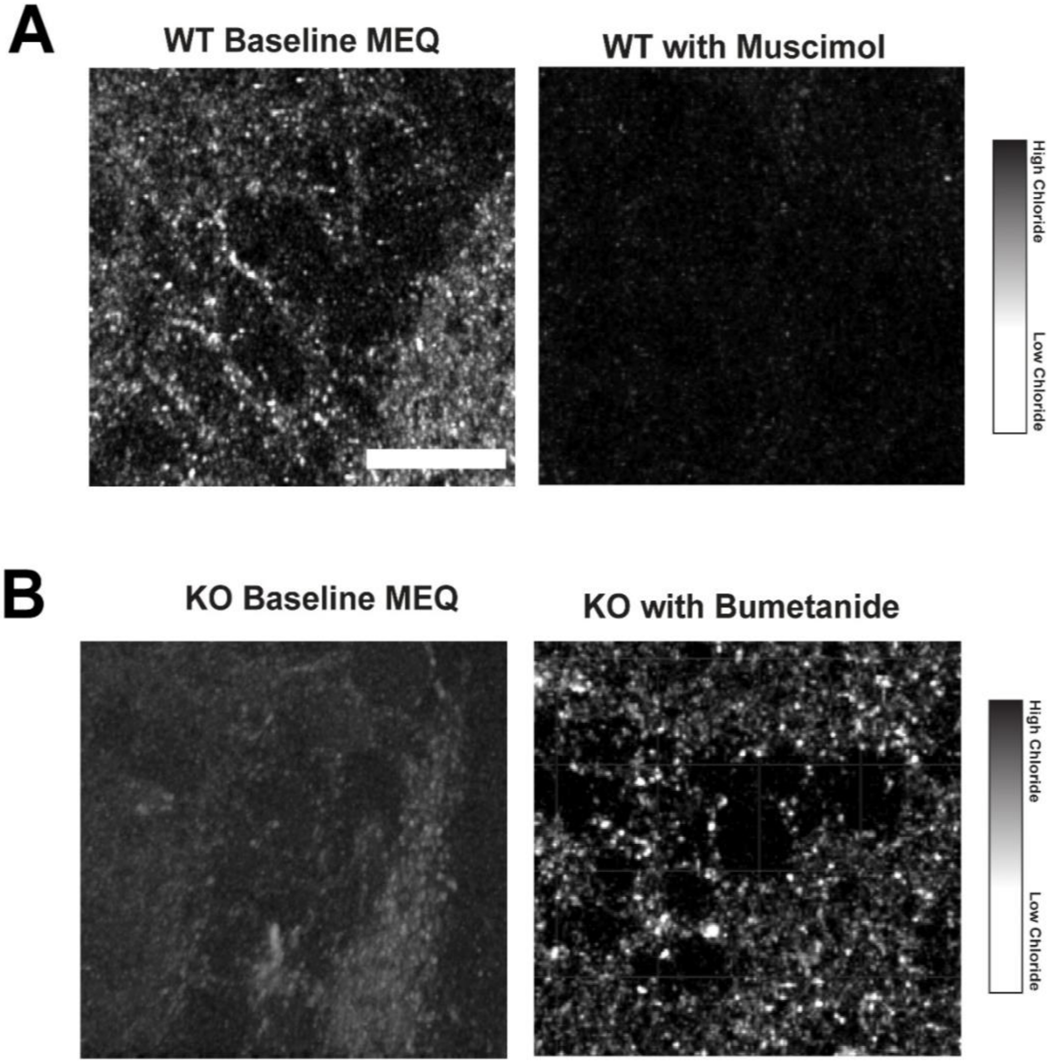


**Supplemental Figure 4*: Controls for Confocal Live Images of Pyramidal Layer for Baseline MEQ Fluorescence.*** *A.) Acute hippocampal slice from WT tissue for baseline MEQ fluorescence and after incubation with GABA$_A$ agonist muscimol to visually reduce fluorescence to confirm MEQ penetration into slice. B) Acute hippocampal slice from KO tissue for baseline MEQ fluorescence and after incubation with bumetanide to visually elevate fluorescence to confirm MEQ penetration into slice*

| **Action Potential Property** | **NHE6 WT**<br><br>**(mean ± SEM)**<br><br>**n=7 cells** | **NHE6 KO**<br><br>**(mean ± SEM)**<br><br>**n=7 cells** | **P-value**<br><br>**(two-tailed, unpaired t-test)** |
|---|---|---|---|
| Onset (mV) | -38.13 ± 2.14 | -43.14 ± 3.54 | 0.2499 |
| Offset (mV) | -39.44 ± 2.83 | -43.23 ± 3.69 | 0.4303 |
| Peak (mV) | 47.05 ± 8.83 | 44.93 ± 3.24 | 0.6566 |
| Spike threshold (mV) | -33.30 ± 2.36 | -35.66 ± 2.89 | 0.5392 |
| Max rise slope (mV/ms) | 152.8 ± 33.67 | 163.3 ± 26.36 | 0.8103 |
| Max fall slope (mV/ms) | -26.09 ± 7.18 | -39.36 ± 6.51 | 0.1962 |
| Duration (ms) | 26.70 ± 4.49 | 38.04 ± 7.78 | 0.1608 |
| *Half width (ms)* | *4.04 ± 0.24* | *2.63 ± 0.16* | **** 0.0004* |
| *Base width (ms)* | *9.043 ± 0.63* | *6.38 ± 0.57* | *** 0.0084* |
| Rise time (ms) | 1.54 ± 0.16 | 1.28 ± 0.12 | 0.2193 |
| *Fall time (ms)* | *7.45 ± 0.53* | *5.08 ± 0.54* | *** 0.0088* |
| Rise rate (mV/ms) | 60.84 ± 13.33 | 69.42 ± 9.77 | 0.6129 |
| *Fall rate (mV/ms)* | *-11.19 ± 1.40* | *-17.20 ± 1.73* | ** 0.0193* |

***Table 1:** **Action potential properties of CA1 pyramidal neurons in response to a 250pA step current injection, average of first 5 action potentials.***

| **Action Potential Property** | **NHE6 WT (mean ± SEM) n=6 cells** | **NHE6 KO (mean ± SEM) n=7 cells** | **P-value (two-tailed, unpaired t-test)** |
|---|---|---|---|
| Onset (mV) | -53.9 ± 4.6 | -56.2 ± 2.2 | 0.6485 |
| Offset (mV) | -52.2 ± 4.4 | -54.7 ± 3.1 | 0.6486 |
| *Peak (mV)* | *18.86 ± 2.8* | *12.4 ± 1.3* | **p=0.0496* |
| Spike threshold (mV) | -34.9 ± 3.2 | -37.2 ± 2.2 | 0.5583 |
| Max rise slope (mV/ms) | 104.5 ± 12.1 | 102.0 ± 7.9 | 0.8584 |
| Max fall slope (mV/ms) | -81.37 ± 11.7 | -83.9 ± 11.04 | 0.8760 |
| Duration (ms) | 14.6 ± 2.2 | 13.2 ± 1.2 | 0.5651 |
| Half width (ms) | 0.94 ± 0.07 | 0.89 ± 0.09 | 0.6507 |
| Base width (ms) | 1.94 ± 0.14 | 1.82 ± 0.19 | 0.6512 |
| Rise time (ms) | 0.96 ± 0.04 | 0.93 ± 0.07 | 0.6846 |
| Fall time (ms) | 0.97 ± 0.11 | 0.89 ± 0.12 | 0.6499 |
| Rise rate (mV/ms) | 57.3 ± 6.7 | 56.1 ± 4.4 | 0.8791 |
| Fall rate (mV/ms) | -60.7 ± 8.9 | -62.6 ± 8.5 | 0.8815 |

***Table 2: Action potential properties of CA1 PV neurons in response to a 250pA step current injection, average of first 5 action potentials. Fluorescence***
